# Gut carriage of antimicrobial resistance genes in women exposed to small-scale poultry farms in rural Uganda: a feasibility study

**DOI:** 10.1101/2020.02.13.947226

**Authors:** Ana A. Weil, Meti D. Debela, Daniel M. Muyanja, Bernard Kakuhikire, Charles Baguma, David R. Bangsberg, Alexander C. Tsai, Peggy S. Lai

**Affiliations:** Department of Medicine, Massachusetts General Hospital Boston, MA, USA; Harvard Medical School, Boston, MA, USA; Mbarara University of Science and Technology, Mbarara, Uganda; Oregon Health & Science University, Portland State University School of Public Health, Portland, OR, USA; Harvard Center for Population and Development Studies, Cambridge, MA, USA; Department of Psychiatry, Massachusetts General Hospital, Boston, MA, USA; Harvard T.H. Chan School of Public Health, Boston, MA, USA

## Abstract

**Background:** Antibiotic use as growth promoters for livestock is presumed to be a major contributor to the acquisition of antimicrobial resistance (AMR) genes in humans, yet data evaluating AMR patterns in the setting of animal exposure are limited to observational studies that do not capture data from prior to livestock introduction.

**Methods:** We performed a feasibility study by recruiting a subset of women in a delayed-start randomized controlled trial of small-scale chicken farming in order to examine the prevalence of clinically-relevant AMR genes. Stool samples were obtained at baseline and one year from five intervention women who received chickens at the start of the study, six control women who did not receive chickens until the end of the study, and from chickens provided to the control group at the end of the study. Stool was screened for 87 clinically significant AMR genes using a commercially available qPCR array (Qiagen).

**Results:** Chickens harbored 23 AMR genes from classes also found in humans as well as vancomycin and additional β-lactamase resistance genes. After one year of exposure to chickens, six new AMR genes were detected in controls and seven new AMR genes were detected in the intervention group. Women who had direct contact with the chickens sampled in the study had greater similarities in AMR resistance gene patterns to chickens than those who did not have direct contact with chickens sampled (p = 0.006). There was a trend towards increased similarity in AMR gene patterns with chickens at one year (p = 0.12).

**Conclusions:** Chickens and humans in this study harbored AMR genes from many antimicrobial classes at both baseline and follow up timepoints. Studies designed to evaluate human AMR genes in the setting of animal exposure should account for high baseline AMR rates, and consider collecting concomitant animal samples, human samples, and environmental samples over time to determine the directionality and source of AMR genes.

Trial registration: ClinicalTrials.gov Identifier: NCT02619227

## INTRODUCTION

Antimicrobial resistance (AMR) is a global public health crisis. Although estimates vary on the severity of the problem, one report has suggested that by 2050, 10 million deaths a year worldwide will be attributed to antimicrobial resistance [1], with crude estimates of the annual economic costs totaling 55 billion dollars in the United States alone [2]. This problem is accentuated in resource-limited settings, where there is a high burden of infectious disease, little to no antimicrobial stewardship, limited resources for microbiology testing, and where it is difficult to obtain access to antibiotics that target highly resistant pathogens. Although prior AMR studies have focused on hospitalized patients and recent administration of antimicrobials to treat infections, an updated view of AMR as a public health problem has highlighted the importance of AMR as a “One Health” problem; that is, viewing human, animal, and environmental health as interconnected and interdependent [3-5]. Antibiotics are widely used in livestock farming to enhance animal health and increase productivity [6], and this practice is thought to be a major contributor to the problem of AMR among humans. However, most available studies are cross-sectional and/or focused on single organisms or pathogens [7-9], and these study designs lack the ability to determine causality. More robust study designs are needed to determine the effect size that antimicrobials used in livestock farms has on transmission of AMR genes to humans.

Surveillance data in 2005 showed that livestock production in Uganda accounted for about 5% of total Ugandan gross domestic product [10], with an estimated annual production of 70.8 million total livestock including cattle, pigs, sheep and goats, and poultry [11]. Studies of poultry farms in Uganda have identified multiple mechanisms of AMR in *Escherichia coli* strains isolated from healthy chickens [12, 13], suggesting that poultry farms may serve as a reservoir of AMR genes for humans. Few studies have evaluated how the initiation of chicken farming relate to AMR in humans, partly due to difficulty in obtaining pre-intervention samples for AMR testing. In this feasibility study, we leveraged an ongoing randomized controlled trial of small-scale poultry farms in rural Uganda to determine patterns of AMR genes in stool of chicken farmers before and after poultry introduction.

## MATERIALS AND METHODS

### Study design and study population

We recruited participants from an existing randomized clinical trial (RCT) of small-scale chicken farming (ClinicalTrials.gov Identifier: NCT02619227) [14]. In this waitlist-controlled RCT conducted in 2015, 92 women living in Mbarara, Uganda were recruited and randomized to receive training, raw materials, and broiler hybrid chicks either immediately (intervention group), or after a 12-month delay (control group). Chicken coops were constructed prior to receipt of the chicks, and study participants were the primary caretakers for the broilers. Broiler chicks were sourced from a single distributor based in Kampala, Uganda and underwent a standard care protocol. Under supervision, participants administered vaccines to the chicks against Newcastle, Gumboro, fowl typhoid, and fowl pox. Participants also routinely administered tetracycline-containing dietary supplements to chicks during the brooding period as part of a protocol to boost growth. Chicken feed was sourced from a single distributor based in Mbarara, Uganda. Routine surveys were administered to monitor behaviors such as recent antimicrobial use (in both chickens and humans) and vaccination status in chickens.

The timing of stool sample collection is depicted in **Figure S1**. Stool samples from six chicken coops belonging to the 6 control participants were collected by retrieving fresh chicken stool once at approximately 18 months after randomization, after the control group had received their chickens as part of the delayed-start randomized controlled trial design. The 6 control participants were chosen based on participants who had stool samples collected from the chickens, and the 5 intervention participants lived in the same villages as the control participants. At baseline, before chickens were introduced into the intervention households and at 12-month follow up after chicken introduction in the intervention group, we obtained fresh stool samples from participants during research clinic visits. The stool samples were frozen within one hour of collection in generator-backed −80°C freezers in the research laboratories of the Mbarara University of Science and Technology. All samples were subsequently transported on dry ice to Massachusetts General Hospital for further processing. All study procedures were approved by the Research Ethics Committee of Mbarara University of Science and Technology (Protocol #30/11-14) and the Partners Human Research Committee (Protocol #2015P000227/BWH). Consistent with national guidelines, we also received clearance for the study from the Ugandan National Council of Science and Technology (Protocol #HS 1746) and the President’s office.

#### Sample processing, AMR gene identification and quantification

Microbial DNA was extracted from 100mg of chicken and human stool samples, and from a reagent-only negative control using the PowerSoil DNA extraction kit (Qiagen, Valencia, CA) according to the manufacturer’s instructions. The presence of AMR genes was screened using a commercially available AMR gene identification microbial DNA polymerase chain reaction (PCR) array (Qiagen, Valencia, CA, cat. No. 330261) according to the manufacturer’s instructions. This array targets six major classes of antibiotics (aminoglycoside, β–lactam, erythromycin, fluoroquinolone, macrolide– lincosamide–streptogramin B, tetracycline, and vancomycin) and includes genes with multi-resistance potential. Briefly, 500ng template microbial DNA was mixed with 1275 µl qPCR mastermix (Qiagen) and nuclease-free water was added to reach a final volume of 2550 µl. 25 µl of reaction mix was added to a 96-well PCR plate containing a pre-dispensed mixture of lyophilized primers and probes for each of the 87 AMR genes. qPCR was performed using Applied Biosystems 7500 Fast Real-Time PCR System using thermal cycling conditions of initial denaturation at 95°C for 10 minutes, followed by 40 cycles of denaturation at 95°C for 15 seconds, and annealing at 60°C for 2 minutes. Raw cycle threshold (CT) values were analyzed using the Microbial DNA qPCR Array data analysis template. One replicate per sample was tested. The efficiency of the PCR instrument and the quality of mastermix were determined by measuring the CT for the control sample between 20 and 24. Validity of the control ensured that potential PCR inhibitors in the sample did not interfere with measurements. A no-template and nuclease-free control were also included to evaluate for the presence of environmental contaminants.

#### Data analysis, visualization and statistical analysis

Determination of detection of AMR genes was performed according to the manufacturer’s (Qiagen) guidelines, described here in brief. The presence or absence of each AMR gene was determined as follows: present if ΔCT > 6, not detected if ΔCT <3, and inconclusive if ΔCT was ≥ 3 and ≤6. To visualize the results of AMR gene presence or absence in each sample, we created a heatmap using the *ggplot2* R package [15]. In order to visualize global patterns of AMR genes over time in the human samples and difference between the chicken samples, we chose to use the Jaccard dissimilarity index. Briefly, the Jaccard index calculates the proportion of unshared features (here AMR genes) out of the total number of features (here AMR genes) recorded between any two samples. To calculate the Jaccard index, we first created a sample by feature matrix denoting the presence or absence of each AMR gene in each sample. Presence/indeterminacy/absence were determined using the ΔCT method described above according to manufacturer recommendations, with the following value assignments; present = 1, indeterminate = 0, absent = 0. Visualization of the dissimilarities in AMR gene patterns was performed using the plot_ordination() function as implemented in the *phyloseq* R package [16]. To evaluate similarity between AMR gene patterns in human samples compared to chicken samples, both at baseline and followup, we computed the distance between the Jaccard index of each sample to the centroid of all chicken samples [17]. In this plot, a shorter distance between data points indicates increased similarity in AMR gene patterns. Measurements were calculated using the dist_between_centroids() function implemented in the *usedist* R package [18]. For statistical testing, we performed a linear regression where the outcome was the calculated distance between each sample and the centroid of the chicken samples, and covariates were group membership (intervention vs control) and time (baseline vs follow-up). All statistical analyses were performed in the R programming language [19]. Two-sided p values of < 0.05 were considered statistically significant.

#### Data Availability

Raw qPCR data results are listed in Supplemental Table 1.

## RESULTS

We collected stool from five women in the intervention group and six women in the control group, from 11 separate households in Nyakabare parish, Mbarara district, Uganda. Mbarara is located in a rural area of Uganda approximately 260km southwest of Kampala, the capital city. The local economy is largely dominated by animal husbandry, petty trading, subsistence agriculture, and supplemental migratory work. Food and water insecurity are common [20-22]. In this study, samples were collected between August 11, 2015 and June 8, 2017. The median age of participants was 35 years, and self-reported demographic data are listed in **Table 1**. All participants were women involved in subsistence farming. At baseline, 10 of the participants reported regular animal contact and a minority reported recent antibiotic use.

**Table 1.** Baseline characteristics of study participants.

|  | Control | Intervention |
| --- | --- | --- |
| n | 6 | 5 |
| Age, years | 40 [34 - 43] | 33 [25 - 40] |
| Farming | 6 (100%) | 5 (100%) |
| Antibiotic use in prior three months |  |  |
| At 0 months | 1 (17%) | 0 (0%) |
| At 12 months | 1 (17%) | 1 (20%) |
| Animal contact | 5 (83%) | 5 (100%) |
| Village chickens<sup>a</sup> | 2 (33%) | 5 (100%) |
| Cows | 2 (33%) | 2 (40%) |
| Goats | 4 (67%) | 4 (80%) |
| Pigs | 1 (17%) | 0 (0%) |
| Dogs | 2 (34%) | 2 (40%) |
| Cats | 2 (34%) | 2 (40%) |

<sup>a</sup>Village chickens refer to free-range chickens that do not receive vaccinations or medications, and do not require an enclosure.

### AMR genes detected

Stool samples from chickens, and from pre- and post-intervention human control and intervention groups were assayed for AMR genes using a validated quantitative polymerase chain reaction (qPCR) assay. All of the no-template controls and positive PCR controls passed the quality control thresholds determined by the manufacturer (**Table S1**). At baseline, the stool of study participants in both control and intervention groups harbored β-lactamase, aminoglycoside, fluoroquinolone, macrolide and tetracycline AMR genes found in the stool (**Table 2**). Seven new AMR genes were detected after one year in the intervention group, and four of these were present in chickens (SHV, SHV[238G240E], QnrS, QnrB-5 group). Six new AMR genes were detected after one year in the control group, and one of these was present in chickens (CTX-M-1 group). Overall, AMR genes were detected from five classes of antimicrobials in humans, and six classes in chickens.

**Table 2.**
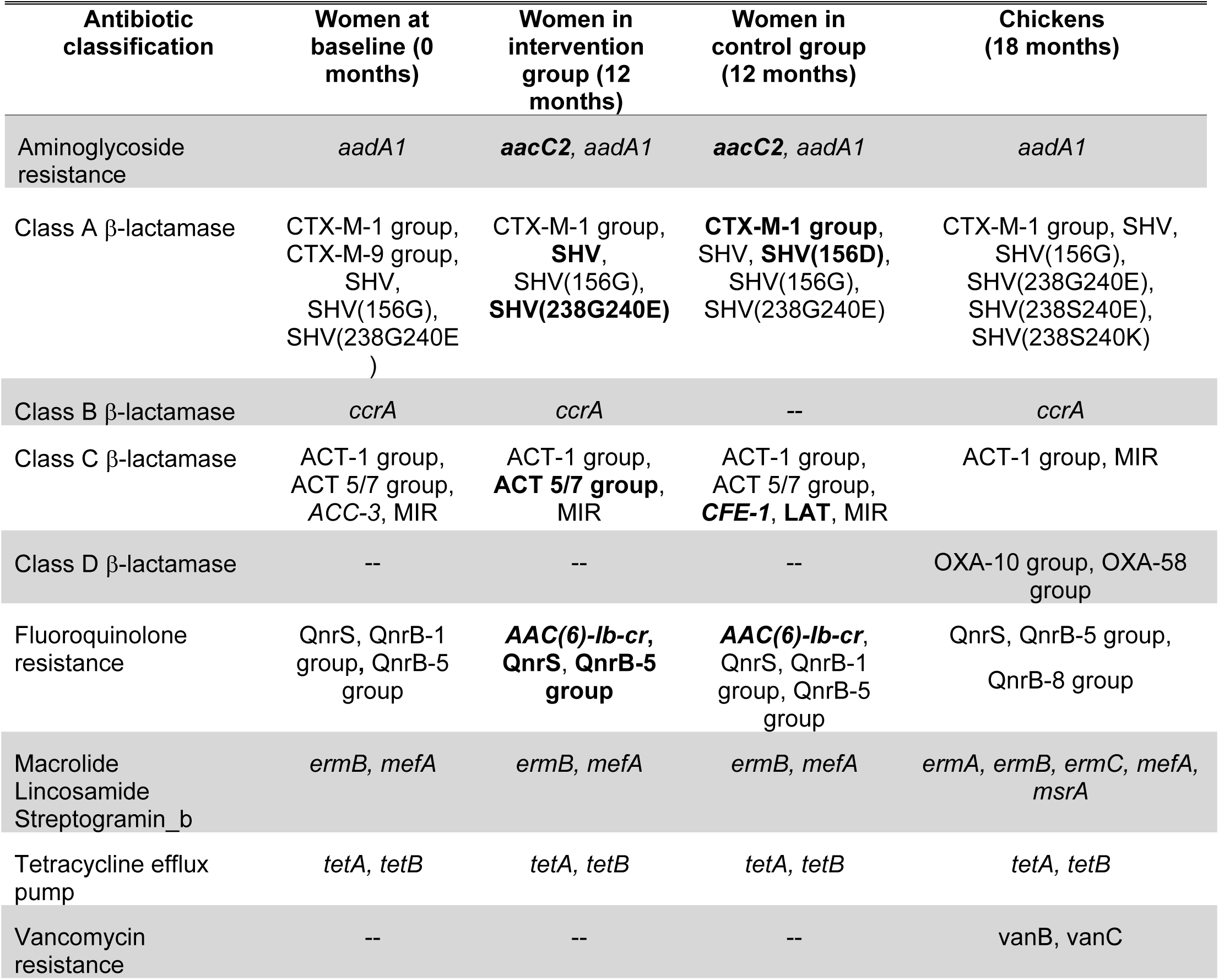
Antimicrobial resistance (AMR) genes in participants detected at baseline and one year post-intervention, and in chickens. Among study participants, newly detected genes after one year are shown in bold. Baseline grouping includes both intervention and control group participants. AMR gene detection was measured using a qPCR array (Qiagen). Raw cycle threshold (CT) values were used to determine detection of AMR, defined as positive if ΔCT >6, not detected if ΔCT <3 and inconclusive if ΔCT was ≥ 3 and ≤6, as per the manufacterer’s instructions. Raw qPCR data is shown in Table S1. Gene names are italicized and names of gene classes are not.

### AMR gene class trends between groups and over time

During the study period there was an overall increase in AMR genes in both the control and intervention groups. The most prevalent AMR genes were *tetA* and *tetB*, which confer tetracycline efflux pumps, and these were found in all chickens tested. *tetA* and *tetB* were also found in the majority of human participants in the study at baseline and follow-up timepoints, as shown in a heatmap of our overall results (**Figure 1** and **Table S1**). β-lactamases were also highly prevalent in both humans and chickens, with Class A and C β-lactamase AMR genes found in humans at both baseline and follow-up timepoints, regardless of chicken exposure, and the Class C β-lactamase MIR present in nearly all study participants. However, Class D β-lactamases were found only in chickens. AMR genes in the Class C β-lactamase group, which includes the clinically important *ampC* β-lactamases responsible for inducible resistance upon exposure to specific antibiotics were particularly dynamic over time, with two AMR genes emerging in the control group after one year that were not seen in other groups (*CFE-1* and LAT), and the loss of *ACC-3*, which was found in the baseline population and not detected upon follow up [23]. Fluoroquinolone and macrolide resistance were widespread over all groups and timepoints. Chicken AMR genes detected included two vancomycin resistance genes that were not found in humans.

**Figure 1.**
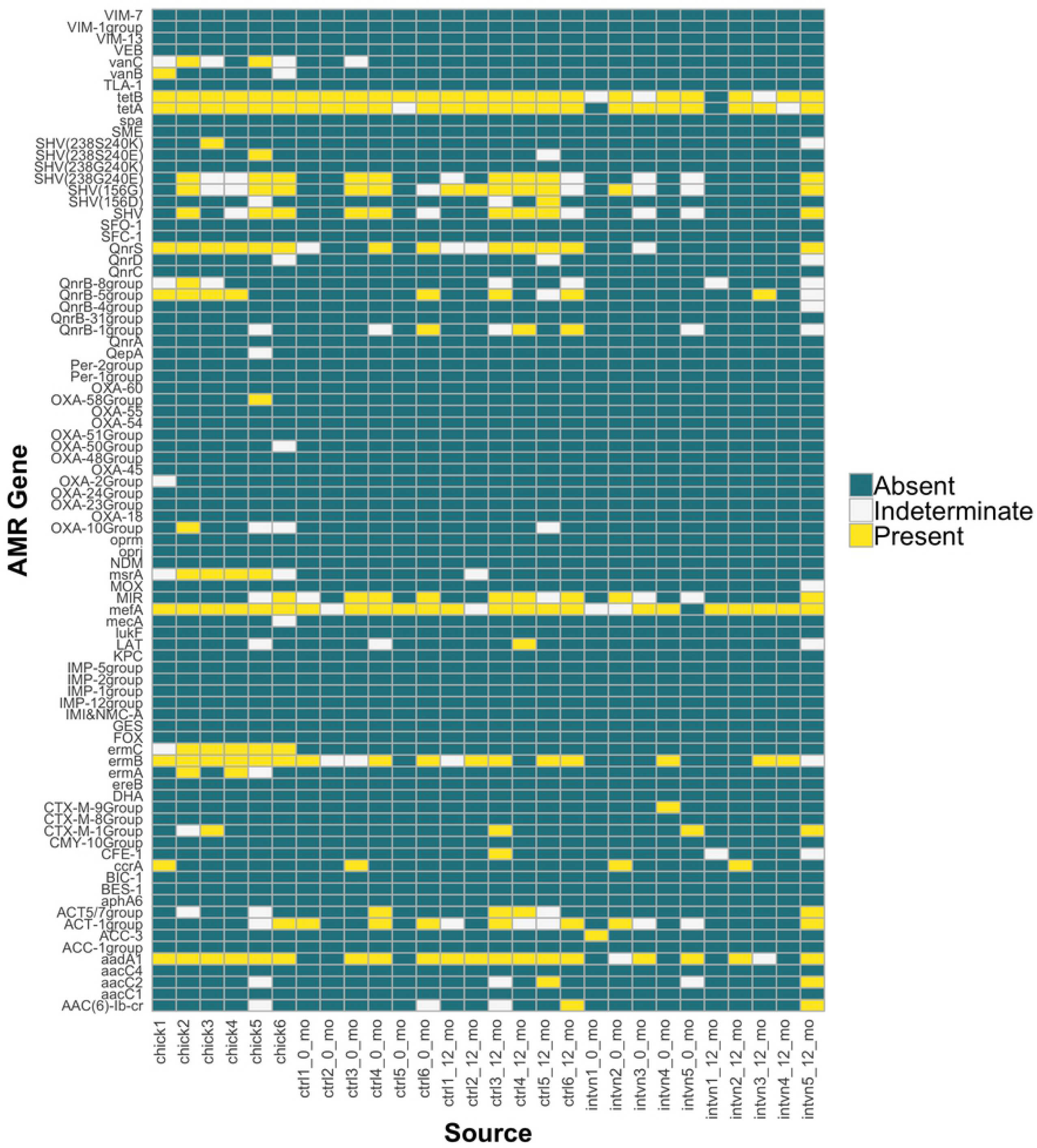
Heatmap demonstrating whether antimicrobial resistance (AMR) genes were present, absent, or indeterminate in human and chicken samples at different timepoints We used an ordination plot to depict patterns of AMR gene composition across groups and over time (**Figure 2)**. To determine the similarity of AMR gene patterns of human samples compared to the chicken samples, both at baseline and follow-up, we computed the distance between the Jaccard index of each sample to the centroid of the chicken samples (**Figure 3**). A shorter distance between data points indicates increased similarity in AMR gene pattern with the chicken samples, while a higher distance indicates decreased similarity in AMR gene pattern with the chicken samples. The AMR gene pattern of the control group is more similar to the AMR gene pattern in their chickens than the intervention group to the control group’s chickens (b = 0.128, p = 0.006, intervention vs. control group; Note more positive b indicates less similarity with chicken samples). There was a trend towards increasing similarity of AMR gene patterns between all humans at one year and chickens though it did not reach statistical significance (b = −0.067, p = 0.12, 12 month vs baseline). In this comparison, the effect size was negative, which is consistent with increased similarity between follow-up AMR gene patterns in both the intervention and control groups compared to chicken AMR gene patterns, though this did not reach statistical significance.

**Figure 2.**
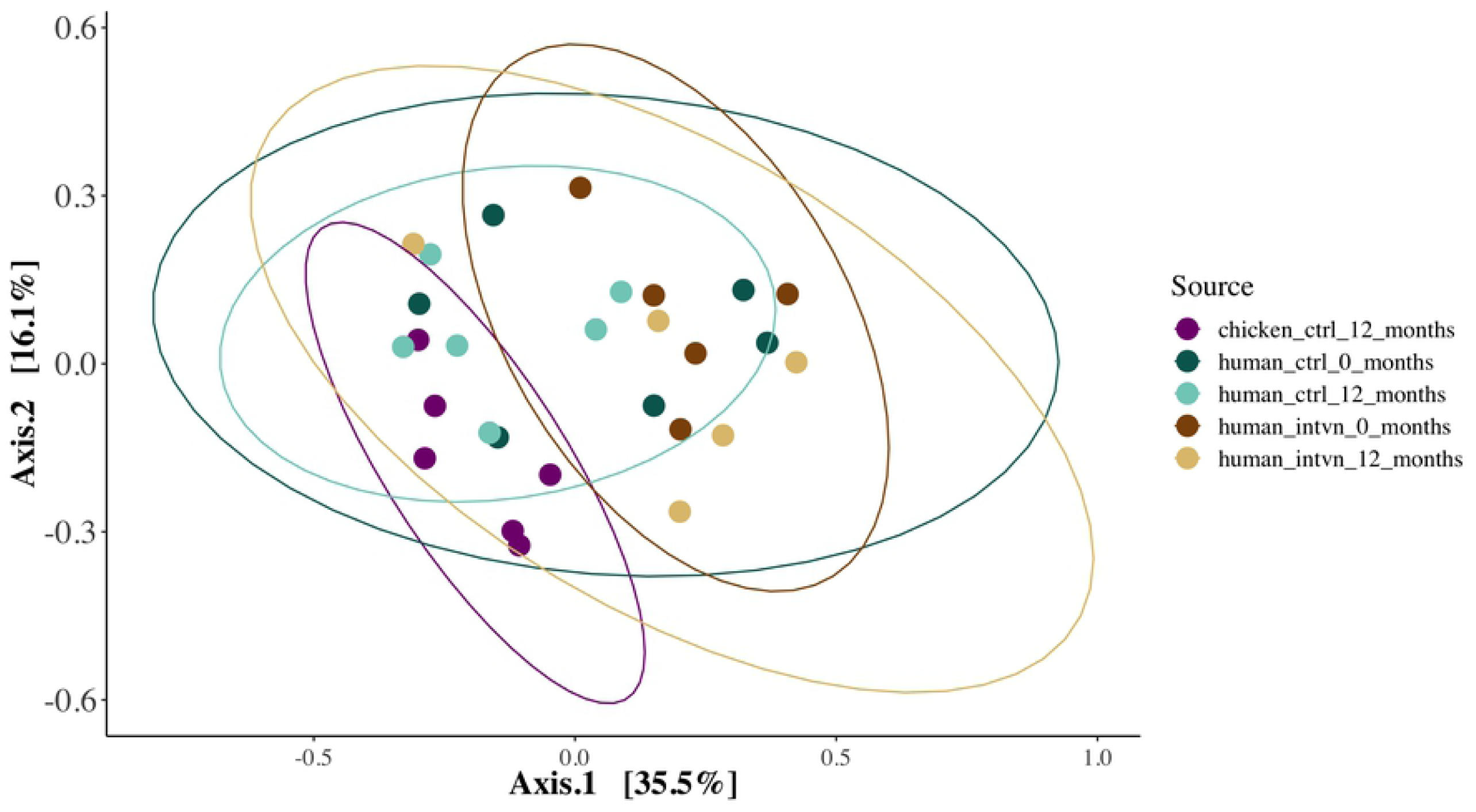
Ordination plot of the Jaccard dissimilarity index of AMR gene patterns between groups. The proportion of unshared AMR genes out of the total number of AMR genes detected between any two samples is shown. More similar samples will appear closer together on the plot. The ellipse depicts the 95% confidence ellipse around each sample group.

**Figure 3.**
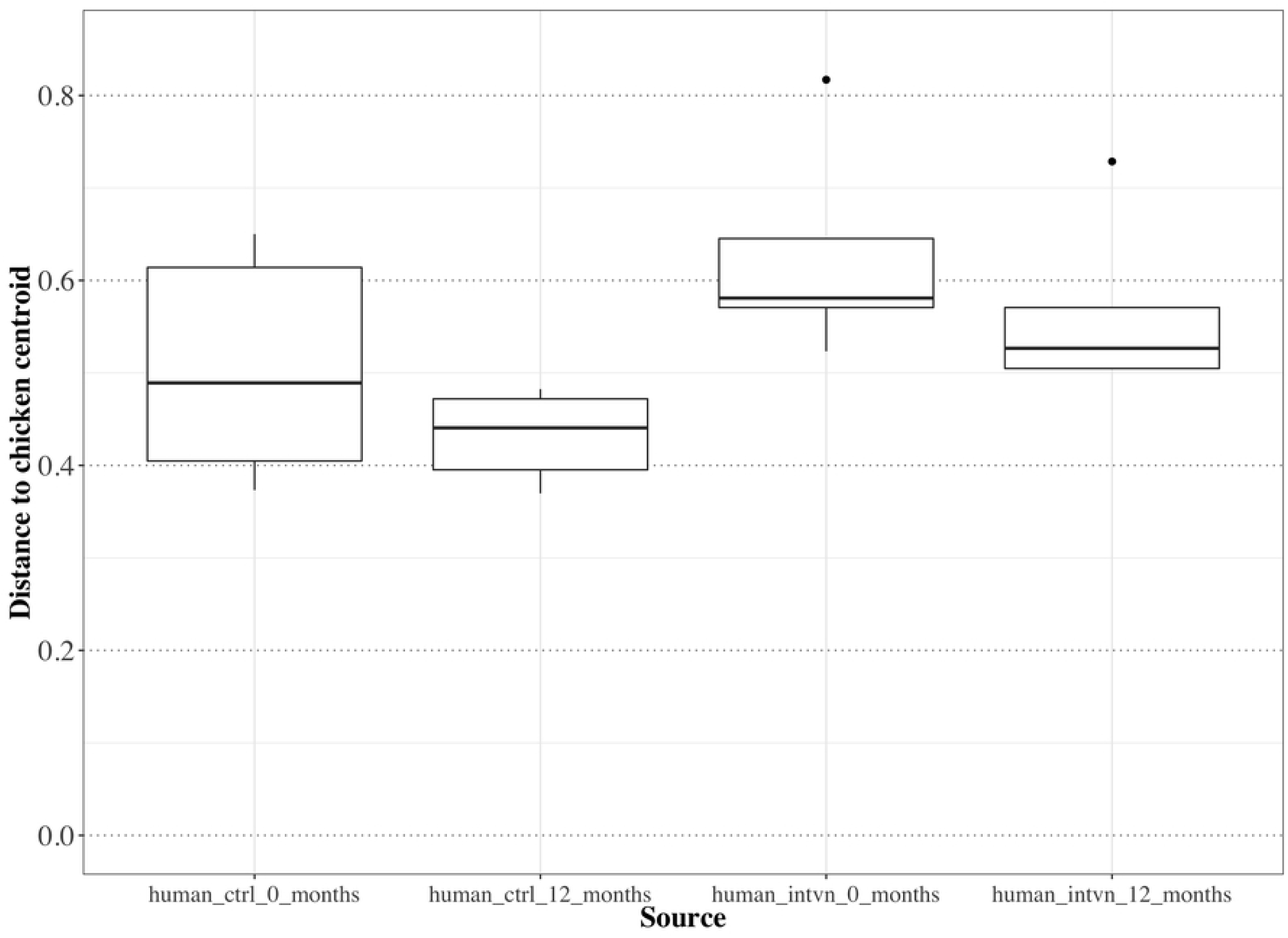
Boxplot of the distance between sample groups and the centroid of the chicken stool samples based on AMR gene pattern. To demonstrate the comparison of the AMR gene pattern of each human sample to the chicken samples at baseline and followup, we computed the distance between the Jaccard index of each sample to the centroid of all chicken samples. Here, a *shorter* distance indicates *increased* similarity in AMR gene pattern of the human sample in relation to the centroid of the chicken samples gene patterns, whereas a *longer* distance indicates *decreased* similarity in AMR gene pattern of that human sample compared to the chicken samples gene patterns. The chicken sample centroid is set at zero. The AMR gene pattern of the chicken samples is more similar to the AMR gene pattern in the control group rather than the intervention group (p = 0.006); note that chicken samples were obtained from the control group. Differences in AMR gene patterns over time did not reach statistical significance (p = 0.12), although at follow-up, the AMR gene patterns in both control and intervention group humans were more similar to AMR gene patterns in chicken samples.

## DISCUSSION

In this study, we find that tetracycline-exposed chickens and humans who care for them harbor AMR genes from multiple gene classes. Over one year, AMR gene carriage increased in all study participants. There were greater AMR gene pattern similarities between humans who had direct interaction with the chickens in the study and the chickens. Over time, AMR gene patterns in both groups became more similar to AMR gene patterns detected in chickens, though this association did not reach statistical significance.

Shared gut organisms between animals and humans increase when close contact occurs between groups, such as in animal husbandry [24, 25]. These shared environments can result in transmission events, which range from zoonotic infections to the spread of benign commensal microbes, or events that represent potential harm to humans, such as acquisition of AMR genes. Pathogens resistant to antibiotics result in more severe illness and increased mortality in humans compared with infections caused by susceptible bacteria [26]. Consistent with the One Health concept, we found that humans and chickens with direct contact had greater similarities in AMR gene carriage in the gut, although the directionality of transmission could not be determined based on our study design. Additionally, we observed a high rate of AMR genes overall in humans and in chickens in rural Uganda.

Tetracycline resistance genes are often found to be widespread among livestock treated with antibiotics, including in Africa [27]. Use of tetracycline in livestock has been associated with increased colony counts of tetracycline-resistant human pathogens in treated animals [28]. While tetracycline was the only antibiotic administered to chickens in this study, a wide range of AMR genes from six different classes were detected in chickens. Many β-lactamase AMR genes with direct links to difficult-to-treat human infections were also detected. For example, the CTX-M-1 Group can confer an extended-spectrum beta lactamase (ESBL) phenotype, and is the most commonly found gene in *Escherichia coli* in the few surveys of AMR genes that have been conducted in African livestock [27]. CTX-M-1 was detected after one year in our control group and was also present in chickens in this study,. CTX-M-1 was also present in the intervention group both at baseline and follow up. While the directionality of CTX-M-1 transmission between control participants and chicken exposure cannot be evaluated in this study, our results demonstrate that CTX-M-1 is circulating in this population among chickens and humans. The carbapenemases OXA-10 and OXA-58 Group were also found in chicken stool in our study and were not found in humans, and confer a concerning degree of antimicrobial resistance [29]. Reasons that these AMR genes have not emerged into the human population are unknown, and may be due to a lack of selective pressure (ie chickens and humans not yet exposed to carbapenem antibiotics) at the time of our study. Similarly, the VanB and VanC genes found in chickens are known to confer vancomycin (glycopeptide antibiotic) resistance to Enterococci, a common genus of the colonic flora, resulting in the clinically important vancomycin resistant enterococcus (VRE). Avoparcin, an antibiotic also from the glycopeptide class, was widely used in livestock and poultry in Europe and linked to VRE isolates in animals. This drug was outlawed for use in animals in the European Union in 1997, although VRE isolates have persisted in some poultry populations after use ceased [30, 31]. Although Avoparcin was not known to be administered to the chickens in this study, it is sold in Uganda as a livestock supplement.

In the humans we studied, numerous AMR genes were acquired over a one year period in both the intervention and control groups. These changes suggest that the population is trending toward increased AMR gene content over relatively short time periods, and that AMR genes in this population are widespread and dynamic. For example, in the fluoroquinolone class, the *AAC(6)-lb-cr* gene is often found on a multiresistance plasmid with other AMR genes, and this gene was detected in both human groups after follow up and was not found in chickens. This indicates that the increased AMR gene patterns over time seen in this population may also originate from sources unrelated to chicken exposure, such as environmental sources [32]. Over one year, we observed that microbial community profiles in humans were significantly altered with (intervention group) or without chicken exposure (control group). We also note shared AMR genes between humans and chickens. Possible explanations for our findings could be that 1) there is a common source of AMR genes in both chickens and humans, for example environmental sources such as water; 2) the possibility exists that AMR genes may be transmitted from humans to chickens; 3) our data does not allow us to comment on transmission of AMR genes from chickens to humans as chicken stool samples were collected after the human stool samples. However, all chicks were from the same distributor and underwent the same care protocol and thus it is possible that some of the AMR genes acquired by humans over time were from direct or indirect chicken contact.

In this study, we describe point prevalence estimates of AMR genes over time. Our study has a few limitations. This pilot study does not evaluate for the directionality or source of transmission of AMR genes detected in humans and chickens as chicken stool samples were not collected simultaneously from all groups at all timepoints. Additionally, our sample size was small, and our detection method often identified gene classes, preventing us from commenting on presence of specific genes. Our qPCR detection of genes were conducted with single replicates due to the cost of the AMR gene arrays. Despite these limitations, our study does highlight the prevalence of circulating AMR genes in people and in chickens in a rural Ugandan population. Our results offer practical design suggestions for future studies evaluating AMR gene transmission in animal husbandry settings. Based on our experience, we would recommend measurement of a wide range of AMR genes at several timepoints, since at baseline a significant number of AMR genes were already present in humans. Sampling from humans, livestock, as well as shared environmental samples (such as water sources) would be required to establish patterns of temporal transmission. A randomized controlled trial design for livestock exposure, as well as molecular evaluation of genetic similarity between bacterial strains harboring AMR genes will be critical for evaluating causality and directionality of transmission.

The World Health Organization Expert Guidelines Development Group tasked with addressing the worldwide crisis of increasing AMR recommend complete restriction of all classes of medically important antibiotics in food-producing animals for growth promotion [33]. Although this was issued as a strong recommendation, evidence to support the recommendation was deemed “low-quality” due to a lack of supportive studies. Here, we describe significant changes in the AMR gene profile in stool of humans over time, and highlight the prevalence of AMR genes in both humans and livestock in a rural Ugandan population. In future studies, to confirm the suspected epidemiologic links that may be responsible for the results in this pilot study, genotyping methods to define mobile elements and strain-specific analysis of AMR genes found in humans exposed to antibiotic-treated livestock are needed. A randomized trial design where simultaneous acquisition of human, livestock and environmental samples may be useful to define susceptibility factors for acquisition of AMR genes.

## SUPPLEMENTARY MATERIAL LEGEND

**Figure S1**: Study design overview.

**Table S1**: Cycle threshold (CT) values of AMR detection from human and chicken stool samples and controls used in this study. PPC=Positive PCR Control.

## ACKNOWLEDGEMENTS

The authors thank the participants in this study. In addition to the named study authors, HopeNet Study team members who contributed to data collection and/or study administration during all or any part of the study were as follows: Phiona Ahereza, Owen Alleluya, Patience Ayebare, Charles Baguma, Patrick Gumisiriza, Clare Kamagara, Allen Kiconco, Viola Kyokunda, Patrick Lukwago, Moran Mbabazi, Juliet Mercy, Elijah Musinguzi, Sarah Nabachwa, Elizabeth Betty Namara, Immaculate Ninsiima, and Mellon Tayebwa. Clean Air Study team members who contributed to data collection and/or study administration during all or any part of the study were as follows: Solome Kobugyenyi, Alex Mutungi, John Bosco Tumuhimbise, Joy Namara Karakire, Collins Agaba. A preliminary analysis of these data was presented at IDWeek, San Francisco, California, USA, October 5, 2018. Funding was provided by National Institutes of Health grants K23 ES023700 (PSL), P30 00002, and K23 MH096620 (ACT), and K08AI123494 (AAW) (https://www.nih.gov/), Harvard School of Public Health-National Institute of Environmental Health Sciences and Center for Environmental Health (P30ES000002) Pilot Project Grant (PSL) (https://www.hsph.harvard.edu/niehs/), American Lung Association Biomedical Research Grant RG-346990 (PSL) (https://www.lung.org/), Harvard Catalyst (UL1 TR001102) Early Clinical Data Support Pilot Grant (PSL) (https://catalyst.harvard.edu/), and Friends of a Healthy Uganda (DRB, ACT) (https://attackpoverty.org/locations/friends-of-uganda/). The funders had no role in study design, data collection, analysis, decision to submit the work for publication, or preparation of the manuscript.

